# Exogenous TGFβ1 and its mimic HpTGM attenuate the heart’s inflammatory response to ischaemic injury and improve long term cardiac outcomes

**DOI:** 10.1101/2023.04.18.537417

**Authors:** Rachael E. Redgrave, Esha Singh, Simon Tual-Chalot, Catherine Park, Darroch Hall, Karim Bennaceur, Danielle J. Smyth, Rick M. Maizels, Ioakim Spyridopoulos, Helen M. Arthur

## Abstract

**Rationale:** Successful and timely coronary reperfusion following acute ST-elevation myocardial infarction (STEMI) is standard therapy to salvage transiently ischaemic heart muscle. However, the subsequent inflammatory response within the infarct can lead to further loss of viable myocardium. Robust interventions are required in the acute MI setting to minimise cardiac injury and reduce risk of further detrimental progression.

**Objective:** TGFβ1 is an anti-inflammatory cytokine released endogenously in response to infection or tissue injury. The goal of this study was to investigate its protective effects when given exogenously following myocardial infarction.

**Methods and Results:** TGFβ1 is found at increased levels in the blood of STEMI patients immediately following myocardial infarction. We observe a significant correlation (p=0.003) between higher circulating TGFβ1 levels at 24h post MI and a reduction in infarct size over the following 3 months, suggesting that an early increase in circulating TGFβ1 is protective in these patients. Using a mouse model of cardiac ischaemia-reperfusion we demonstrate that additional exogenous TGFβ1 delivered in the acute setting has multiple beneficial outcomes. At 24 hours post-reperfusion It leads to a significantly smaller infarct size (30% reduction, p=0.025), reduced inflammatory infiltrate (28% reduction, p=0.015), lower intra-cardiac expression of inflammatory cytokines IL1β and CCL2 (>50 % reduction, p=0.038 and 0.0004, respectively) and reduced scar size at 4 weeks (21% reduction, p=0.015). Furthermore exogenous delivery of an equivalent dose of HpTGM, a recently described low-fibrogenic mimic of TGFβ1, secreted by a helminth parasite to evade immune rejection, has an almost identical protective effect on injured mouse hearts. Furthermore using a genetic approach we show the benefit is mediated by the vascular endothelium.

**Conclusions:** This work reveals the potential of exogenous TGFβ1 and HpTGM delivered in the acute MI setting to provide protective anti-inflammatory effects and reduce infarct size, leading to a smaller scar and reduced detrimental progression.

## Introduction

Despite major improvements using primary percutaneous intervention (PPCI) to treat patients with acute ST elevation myocardial infarction (STEMI), progression to heart failure post-infarction represents a major clinical problem^1, 2^. It has been reported that 22% STEMI patients treated with PPCI progressed to develop heart failure symptoms within 1 year, despite state-of-the-art medical care^3^. Detrimental progression is substantively determined by the original infarct size and time to reperfusion. An acute exuberant pro-inflammatory response can further enhance local cardiac injury leading over time to adverse ventricular remodelling and gradual loss of cardiac function that results in heart failure. For STEMI patients, particularly those with large infarcts, additional intervention in the acute setting is needed to reduce I/R injury, protect myocardial tissue and thereby reduce the risk of progression to heart failure.

The immediate effect of an acute coronary occlusion is cardiomyocyte death due to anoxia. Timely reperfusion is the most effective treatment to save ischaemic myocardium. However, reperfusion itself causes release of damaging reactive oxygen species, whilst necrotic cardiomyocytes release alarmins that activate innate immune cells. Pro-inflammatory cytokines are upregulated within injured myocardium and an enhanced pro-inflammatory environment in the myocardium leads to further increases in immune cell infiltration associated with increased myocardial cell death^4, 5^. Therefore, dampening the immediate pro-inflammatory response following ischaemia-reperfusion has the potential to protect the surviving myocardium from further injury.

TGFβ1 is an endogenous circulating and tissue-resident protein that has multiple critical roles in regulating the development and maintenance of the cardiovasculature^6^. It is stored in a latent form within tissues and is activated by complex proteolytic mechanisms in response to tissue injury^7^. Active TGFβ1 ligand signals via a tetrameric TGFβ receptor protein complex on the cell surface that activates SMAD2/3 transcription factors to regulate gene expression^8^. The pleiotropic properties of TGFβ1include being a major driver of anti-inflammatory responses ^9-12^. In line with this property, mice lacking TGFβ1, which survive to birth, die in early post-natal life from multifocal inflammatory autoimmune disease ^13, 14^. Furthermore, in a mouse model of atherosclerosis, inhibition of TGFβ signalling promotes development of atherosclerotic lesions with an increased inflammatory component^15^. In injured heart tissue, following MI, TGFβ1 is critical for promoting the transition from an early pro-inflammatory phase to a later pro-reparative phase of cardiac healing. It drives this transition via a number of mechanisms including promoting the formation of anti-inflammatory T regulatory cells (Tregs), stimulating angiogenesis and initiating fibrosis^7, 16^. It has additional anti-inflammatory effects on the vascular endothelium, including reducing expression of e-selectin, required for leukocyte adhesion prior to extravasation into the injured tissue^17, 18^. An early trigger of some of these anti-inflammatory properties is expected to be beneficial in dampening an exuberant pro-inflammatory response immediately post-infarction. In line with this hypothesis, using rat and feline models of cardiac ischaemia reperfusion, treatment with exogenous TGFβ1 in the acute setting (i.e. during ischaemia and prior to reperfusion) reduces myocardial injury within the first 24hours of reperfusion,^19 20^. However, these studies did not investigate the more important longer-term effects on cardiac outcomes. In this study we confirm the myocardial protective role of exogenous TGFβ1 delivered in the acute setting of cardiac ischaemia-reperfusion in a mouse model, we show that this involves an early anti-inflammatory effect and leads to reduced scar size at 4 weeks after the initial injury. As TGFβ1 also has robust pro-fibrotic properties ^21^ and therefore unlikely to gain broad favour as a therapeutic agent, we investigated whether its parasite derived mimic HpTGM has similar protective properties. HpTGM is one of a number of proteins secreted by the helminth worm *Heligmosomoides polygyrus* in order to evade immune rejection from the murine gastrointestinal tract. HpTGM has no molecular homology to TGFβ1 but signals through the same TGFβ receptor complex and, importantly for our study, has greatly reduced pro-fibrotic properties compared with TGFβ1 ^22^. Here we report that a single bolus of HpTGM given at the clinically relevant time of coronary artery reperfusion has similar anti-inflammatory protective effects to TGFβ1 leading to reduced scar size.

## Methods

### Patient population

STEMI patients (n=52) were from a previously published double-blind, randomised controlled trial (Cyclosporin to Reduce Reperfusion Injury in Primary PCI Clinical Trial; “CAPRI”)^23^ with CMR imaging available at 1 week and 3months post-MI. The study complied with the principles of the declaration of Helsinki, was approved by the local ethics committee (NCT02390674) and written informed consent was obtained from all participants. Patient details are summarised in Table S1. Patient blood samples were taken at 15 minutes post reperfusion from the culprit coronary artery, and at 24 hours post-reperfusion from a peripheral vein. Serum was analysed using a Luminex-based custom xMAP immunoassay (BioRad) with TGFβ1 detection beads.

### Mouse surgery

All animal experiments were performed under EU legislation and approved by the animal ethics committees of Newcastle University. The study used male mice aged between 12-16 weeks of the C57BL/6 strain purchased from Charles River UK. Mice with tamoxifen-induced endothelial-specific depletion of the TGFβ type II receptor (Tgfbr2fl/fl;Cdh5(BAC)-Cre-ert2) have been previously described^24^. Tamoxifen treatment (2mg/day) was given in adults (age 10-12 weeks) by intraperitoneal injection for 5 consecutive days and 14 days prior to surgery. Surgery to induce a 60 minute transient occlusion of left anterior descending (LAD) artery for 60 minutes was performed under isoflurane anaesthetic as previously described^25, 26^, except that pre-surgical sedative/analgesia was provided by subcutaneous injection of midazolam (5mg/Kg) and fentanyl (0.05mg/Kg). Sham controls had the same surgery but without coronary artery ligation. All animals entering the study were subject to the same exclusion criteria as follows: failure to recover from surgery within 2 hours; small infarct as judged by extent (< 30%) of blanched left ventricular cardiac tissue at the time of ligation, no ST elevation observed on ECG during ischaemic period, failure of the occluded artery to reperfuse after 60 minutes, failed injection of drug.

### Drug Treatments

A catheter was placed in the tail vein immediately prior to surgery to enable time-controlled drug delivery. In the first study recombinant murine TGFβ1 (eBioscience #14-8342-82) was given at two time points per mouse 60 minutes apart; the first dose (33ug/kg) was given within one minute of ligation of the LAD artery and the second dose (also 33ug/kg) up to one minute after reperfusion of the LAD. Control mice had the same surgery but no injection. The second study used a cytokine produced by *Heligmosomoides polygyrus* TGM (HpTGM), expressed as a recombinant protein in HEK293T cells and purified using affinity chromatography, as previously described ^22^. A single bolus of HpTGM or an equivalent volume (∼50ul) of saline was given within one minute of reperfusion unless otherwise stated. Of note, the bioactive form of TGFβ1 is a dimer of 25kd, whereas HpTGM is a monomer of 49kd so we used twice the total concentration of HpTGM (132ug/kg). The surgeon was blinded to saline or HpTGM treatment.

### Area of infarct and density of infiltrate

Hearts were harvested at 24 hours post reperfusion and heart cryosections (10 μm) were processed for immunofluorescent staining as previously described ^26, 27^. All staining analyses were blinded to treatment group. Leukocytes were detected using rat primary antibody anti-CD45 (Biolegend #103102), with detection using secondary anti-rat antibodies conjugated to Alexa568 fluor. A secondary antibody only control was used in each case to ensure immunostaining specificity. Immunostaining for leukocytes was used to characterise the density and area of immune cell infiltrate (Figure S1).

To evaluate the extent of non-viable myocardial tissue, the heart was harvested 24 hours after reperfusion, briefly cooled and cut into 6 to 8 slices per heart. The rings were stained in 1% triphenyltetrazolium chloride (TTC) in 0.9% NaCl at 37°C for 20 minutes and fixed in 10% buffered formalin at RT for 1hour. ImageJ analysis of images (taken using an MZ6 Leica microscope) of the TTC-negative (non-metabolically active) tissue compared with the total left ventricular myocardium was used to calculate percent infarction. To ascertain mature scar size at 4 weeks post reperfusion, 10um sections every 20^th^ section from ligature to apex through the heart were stained using the Masson’s Trichrome Stain Kit (Sigma) and imaged using an Aperio slide scanner. Scar size was presented as % total left ventricle.

### Intra-cardiac cytokine expression

Hearts were harvested 24 hours after reperfusion. Left ventricles were dissected, washed in PBS to remove blood, harvested into RNAlater (Thermofisher) and frozen (−80°C) until required. Tissue was finely minced and RNA extracted using the RNeasy fibrous tissue mini kit (Qiagen) according to manufacturer’s instructions. Random hexamers were used to prime cDNA synthesis using the Tetro cDNA synthesis kit (Merdian Bioscience). Cytokine expression was measured using TaqMan Universal PCR master mix and the following Taqman probes: interleukin 1β (IL1β) Mm00434228_m1; chemokine C-C motif ligand 2 (CCL2, also known as MCP1) Mm00441242_m1, tumour necrosis factor (TNFα) Mm00443258_m1 and Tgfb1 Mm01178820. Taqman probes for housekeeping genes were Glyceraldehyde-3-phosphate dehydrogenase (Gapdh) Mm99999915_g1 and hypoxanthine guanine phosphoribosyl transferase (HPRT) Mm03024075_m1. Gene expression was measured in I/R hearts from TGFβ1 or HpTGM treated mice versus naïve hearts using the delta delta Ct comparison method.

### Cell Culture and Western Blot

Cells were derived from mice of the C57BL/6 background strain carrying the floxed *Tgfbr2* allele, tamoxifen inducible *Rosa26-CreERT2* transgene and the temperature sensitive Immorto gene^28^. Mouse coronary endothelial cells (MCECs) were isolated from heart tissue using anti-CD31 conjugated magnetic beads. MCECs were cultured in a similar way to our mouse lung endothelial cell lines ^29-31^ in Promocell MV2 media with 5% serum. To deplete the *Tgfbr2* allele, cells were treated with 1μM 4-hydroxytamoxifen for 48 hours to activate Cre-ERT2 and generate *Tgfbr2 knockout(KO)* MCECs. Cells were cultured in fresh media in the absence of tamoxifen for at least a further 48 hours prior to use. Western blot experiments were performed as previously described^30^ using mouse anti-pSMAD2 (cell signalling #3108S diluted 1/500) detected using anti-mouse IRDye 800CW (Licor #925-32211) and α-tubulin (Sigma #T6199 diluted 1/2000) detected using anti-mouse IRDye 680RD (Licor #925-68070). Fluorescent gel blots were imaged using an Odyssey scanner.

### Statistical Analysis

In the STEMI patient cohort, the primary endpoint was a change in infarct size at 3 months versus 1 week post-PPCI. Spearman correlation analysis was used to determine the association of TGFβ1 levels at 24h with change in infarct size over the 3 month study period. The number of biological replicates is provided in the figure legend. Data were analysed using Graphpad Prism and differences were considered significant where p<0.05. Data were tested for normality using the Shapiro-Wilk test, where n numbers were sufficient, and used to inform the choice of parametric or non-parametric statistical tests. Normally distributed data are presented as mean ± standard error. Students unpaired two-tailed students t test or Mann Whitney test were used to compare two experimental groups, whereas data from multiple groups were compared using one way ANOVA (with Tukey post hoc corrections for multiple comparisons) or Kruskal-Wallis test (with Dunn’s correction). The statistical test used is indicated in the figure legends and significance was set at p<0.05.

## Results

### Circulating TGFβ1 levels at 24h post reperfusion in STEMI patients correlates with beneficial reduction in infarct size after 3 months

In light of the previously reported protective role of TGFβ1 following myocardial infarction in preclinical studies ^19, 20^and established role of TGFβ1 in dampening the inflammatory response ^9-11^ we examined circulating TGFβ1 levels in STEMI patients. We used data from our recent CAPRI trial where early blood samples and 1 week plus 3 month MR cardiac function outcome measures were available^23^. This trial was originally designed to detect whether treatment of STEMI patients with a single bolus of cyclosporin immediately before PPCI reduces the amount of damage to the heart compared to treatment with placebo. We first assessed whether cyclosporin treatment affected serum TGFβ1 levels at the two timepoints tested: 15minutes and 24 hours after reperfusion. No differences in TGFβ1 levels nor infarct size were observed when comparing STEMI patients with respect to their treatment with cyclosporin or placebo (Table S2). We therefore pooled the data for analysis. The median TGFβ1 level was 21,932 pg/ml at 15 minutes post reperfusion and dropped to 6,248 pg/ml at 24 hours, a fall of 3.5-fold (p<0.0001) (Figure 1A). Over the subsequent 3 months under standard optimal medical care, the infarct size of these STEMI patients decreases on average by 2.5% (p<0.01) (Figure 1B). Interestingly, circulating TGFβ1 levels at 24h correlates with the size of this reduction in infarct size (p=0.003; Figure 1C), suggesting TGFβ1 might have beneficial effects.

**Figure 1.**
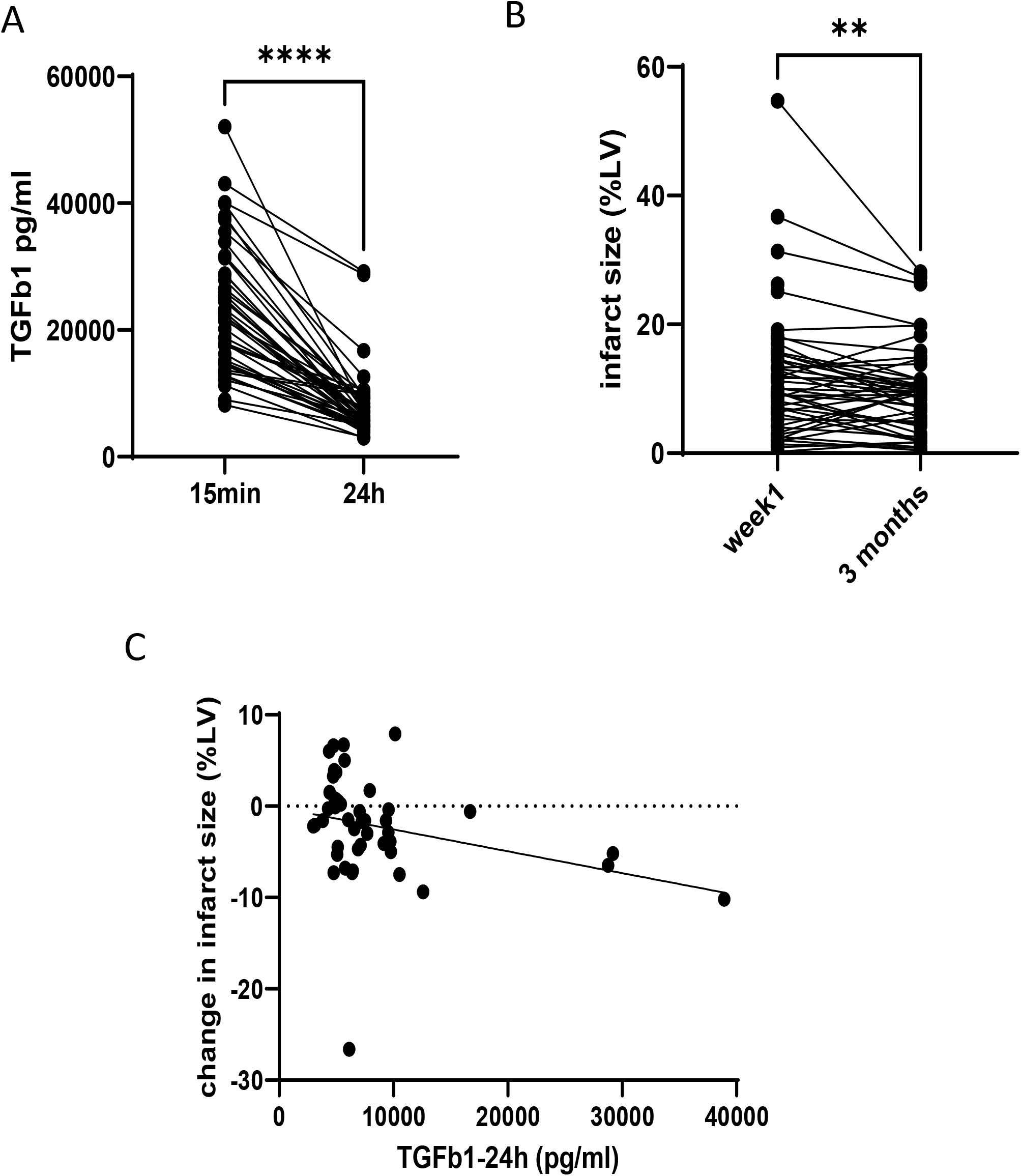
Circulating TGFβ1 levels in STEMI patients at 24h post reperfusion positively correlate with a reduction in infarct size over 3 months. **A**. High circulating levels of TGFβ1 at 15 minutes post reperfusion in STEMI patients fall by an average of 2.7 fold by 24h (p<0.0001, Wilcoxon matched paired samples test). **B**. Infarct size decreases by an average of 2.5% between week1 and 3 months following PPCI in STEMI patients (p=0.0085, Wilcoxon two tailed matched paired signed rank test) **C**. There is a significant inverse correlation between circulating TGFβ1 levels at 24 hours post-reperfusion and change in infarct size over the subsequent 3months. Spearman’s rank correlation coefficient =-0.42; p= 0.0034.

### Exogenous TGFβ1 reduces infarct size and inflammatory responses at 24h post reperfusion

To investigate the effect of TGFβ1 on pathological changes in heart tissue following myocardial infarction, we used our established surgical mouse model of cardiac ischaemia-reperfusion^25, 26^. First we showed that intravenous delivery of recombinant TGFβ1 at the acute phase of ischaemia/reperfusion reduced infarct size at 24h by 30% (p=0.025), based on cell viability staining with triphenyl tetrazolium chloride (TTC) (Figure 2A,B). This TGFβ1-mediated protection agrees with a previous study using a cat model^19^. Cardiac veins are the primary site of leukocyte adhesion and extravasation post reperfusion. We have previously shown that our mouse model of ischaemia reperfusion leads to significant endothelial cell leukocyte adhesion to the venous endothelium by 2h post-reperfusion and there is major leukocyte infiltration of the injured left ventricular tissue by 24 hours ^26^. We therefore analysed the effects of increased acute circulating TGFβ1 levels on the intramyocardial inflammatory infiltrate at 24 hours post-reperfusion using heart tissue immunostained for the pan-leukocyte marker CD45. We found that TGFβ1 treatment leads to a significant reduction of 23% (p=0.03) in the area of leukocyte infiltrate compared with untreated ischaemic hearts (Figure 2 C,D). This area of infiltrate can also be used as an indirect readout of infarct size (Figure S1), in place of the TTC assay. Importantly, the mean density of CD45+ leukocytes within the injured myocardium was also significantly reduced by 28% (p=0.015) in TGFβ1-treated versus untreated mice (Figure 2E). Inflammatory cytokines are known to be released rapidly within the heart following tissue injury. We therefore examined intra-cardiac expression of three key pro-inflammatory mediators: tumour necrosis factor alpha (TNFα), chemokine (C-C motif) ligand 2 (CCL2, also known as MCP1) and interleukin 1β (IL1β). These cytokines were all significantly upregulated in the injured left ventricle (LV) at 24h post reperfusion compared with sham controls (Figure 3). In addition, the expression of CCL2 and IL1b was reduced by over 50% (p=0.004 and p=0.038 respectively) in the injured LV of TGFβ1-treated compared with untreated mice (Figure 3A,B). On the other hand, induced expression of TNFα was relatively modest and was similar in all infarcted hearts (Figure 3C).

**Figure 2.**
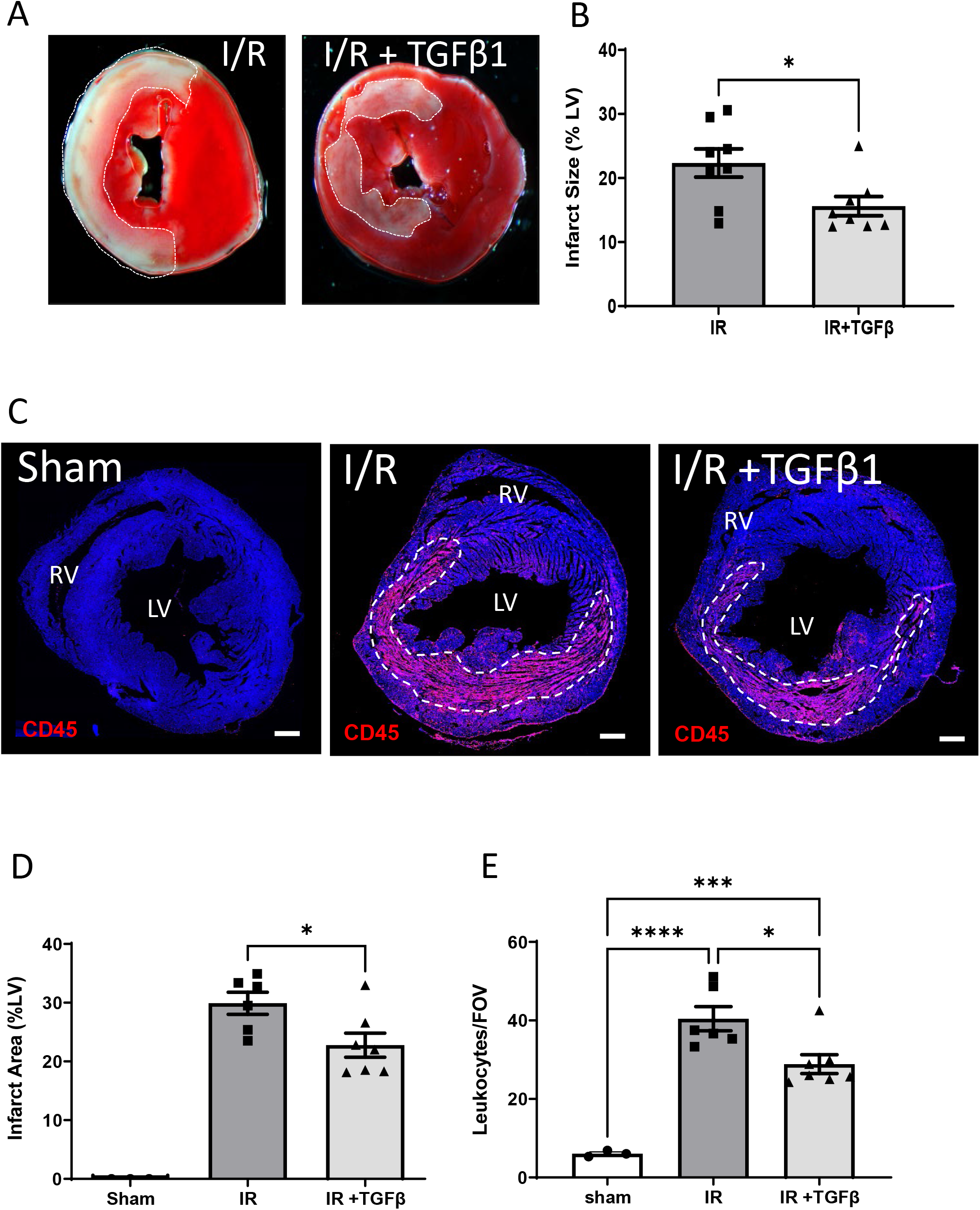
Treatment with exogenous TGFβ1 in a mouse model of transient cardiac ischaemia reduces infarct size and inflammatory infiltrate 24h after reperfusion. **A**,**B** hearts were subject to 60 minutes infarction by transient occlusion of the LAD followed by reperfusion with or without TGFβ1 treatment (66ug/kg). Area of infarct as a proportion of the left ventricle was measured using viable (TTC) staining at 24h post reperfusion. Infarct area of non-viable myocardium (white) as a proportion of the total left ventricle was significantly reduced in the TGFβ1 treated mice (*p<0.05; n=8 per group). **C** Immunostaining of transverse heart sections with anti-CD45 (pan-leucocyte marker) at 24 hours following reperfusion was used to measure the total area of immune infiltrate (dashed line) expressed as % LV. This was then used as a readout of infarct injury and this area of infiltrate is absent in sham controls. **D** TGFβ1 treatment leads to significant reduction of the injured area. Mean values shown for 8 images per biological replicate, taken from transverse heart sections (200um apart) from the intramyocardial region of the mid left ventricle. Data analysed by unpaired two tailed t test (data values in shams were all zero). *p<0.05 n=6-7/group **E** Anti-CD45 staining was used to measure the density of leukocyte accumulation in the injured region at 24 hours post-reperfusion using the quantification methods detailed in supplementary figure 1. TGFβ1 treatment leads to significantly reduced density of CD45+ leukocytes in the injured area. Leukocytes found within the myocardium of sham hearts correspond to tissue resident leukocytes. Data analysed by one way ANOVA with tukey correction. *p<0.05; ***p<0.001; ****p<0.0001; n=6-7/group. Abbreviations: lv left ventricle; rv right ventricle; FOV field of view.

**Figure 3.**
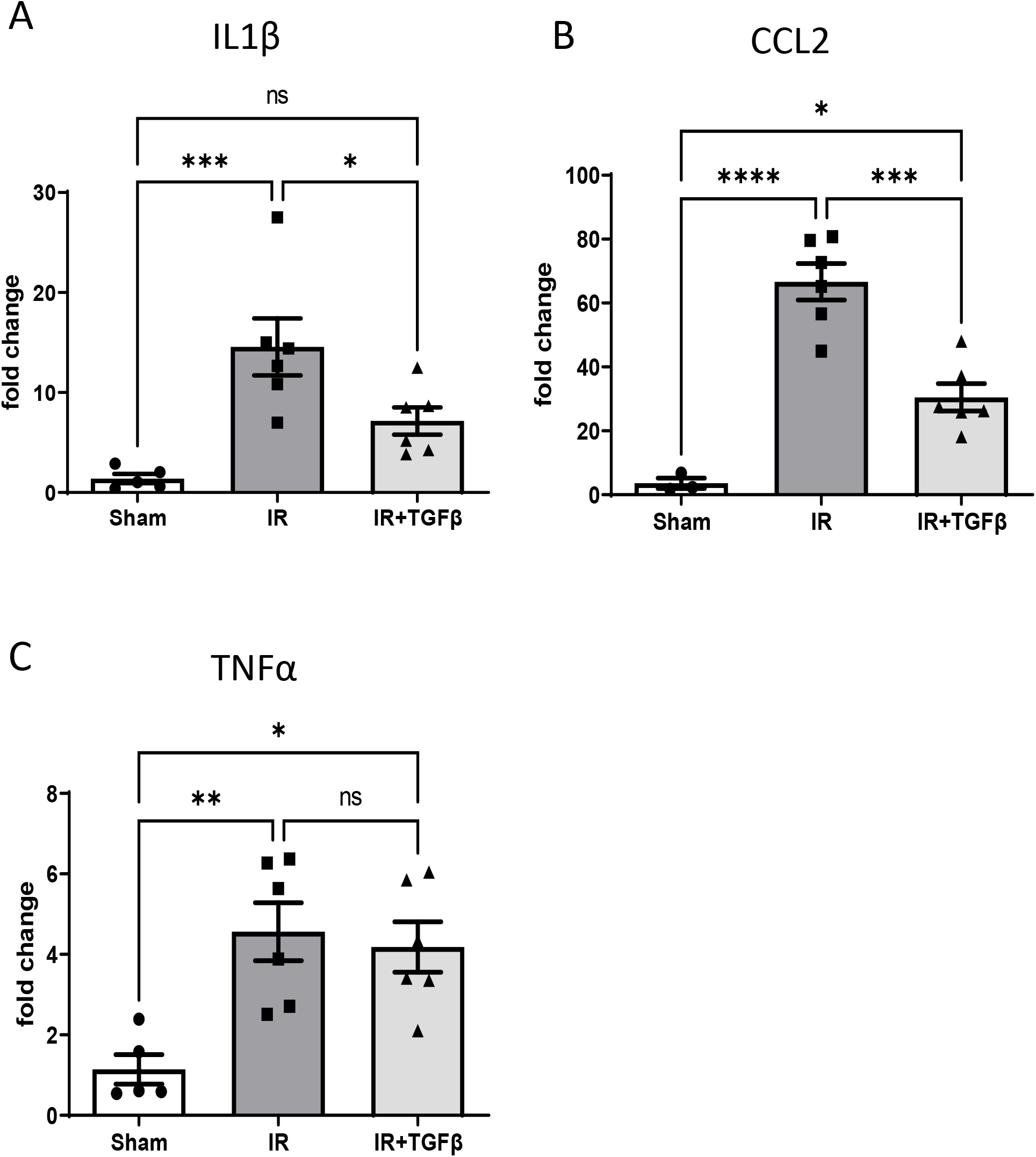
Exogenous TGFβ1 treatment reduces myocardial induced expression of major inflammatory cytokines. **A**,**B**,**C:** qPCR analysis of infarcted (60min) left ventricular heart tissue at 24h post reperfusion shows upregulation of cytokines IL1β, CCL2, and TNFα compared with sham operated hearts. Both IL1β and CCL2 are significantly increased in IR hearts from control mice compared with TGFβ1 treated mice. Gene expression data was normalised to housekeeping genes Hprt1 and Gapdh, and fold change in gene expression was calculated with respect to expression in naïve hearts. Data were analysed by one way ANOVA with tukey correction for multiple comparisons *p<0.05; **p<.01;***p<0.001;****p<0.0001.

### Exogenous TGFβ1 treatment reduces mature scar size at 4 weeks post reperfusion

In our mouse model, transient cardiac ischaemia due to ligation of the LAD for 60 minutes followed by reperfusion leads to a mature intramural collagenous scar by 4 weeks (Figure 4A). Masson’s trichrome staining discriminates viable muscle from collagenous scar and was used to measure mature scar size at 4 weeks post infarction. This intramural scar contrasts with the transmural scars that result from permanent LAD occlusion used in many other mouse studies in the literature, including our own ^32^. The mean scar size is significantly reduced by 21% in the TGFβ1 treated mice compared with untreated mice (p=0.015; Figure 4B). This indicates a longer-term improved outcome and is consistent with the beneficial effects of acute TGFβ1 treatment on infarct size seen at 24h post-reperfusion. Delivery of exogenous TGFβ1 polypeptide has been shown to have an immediate half-life of 11 mins and a terminal half-life of 60 mins^33^. Therefore, the majority of the ligand will have cleared within 24h. Despite these labile properties and the observed benefit of TGFβ1 in *reducing* scar size there are still likely to be concerns around the pro-fibrotic properties of exogenously delivered bioactive TGFβ1 that might prevent its consideration as a therapeutic in STEMI patients. We therefore turned to HpTGM, an immunomodulatory mimic of TGFβ cytokine that is produced by the helminth worm *Heligmosomoides polygyrus* ^22, 34^. HpTGM signals through the TGFβ receptor complex (Figure S2) and mediates similar anti-inflammatory effects whilst lacking the robust pro-fibrotic properties of TGFβ1 ^22^.

**Figure 4.**
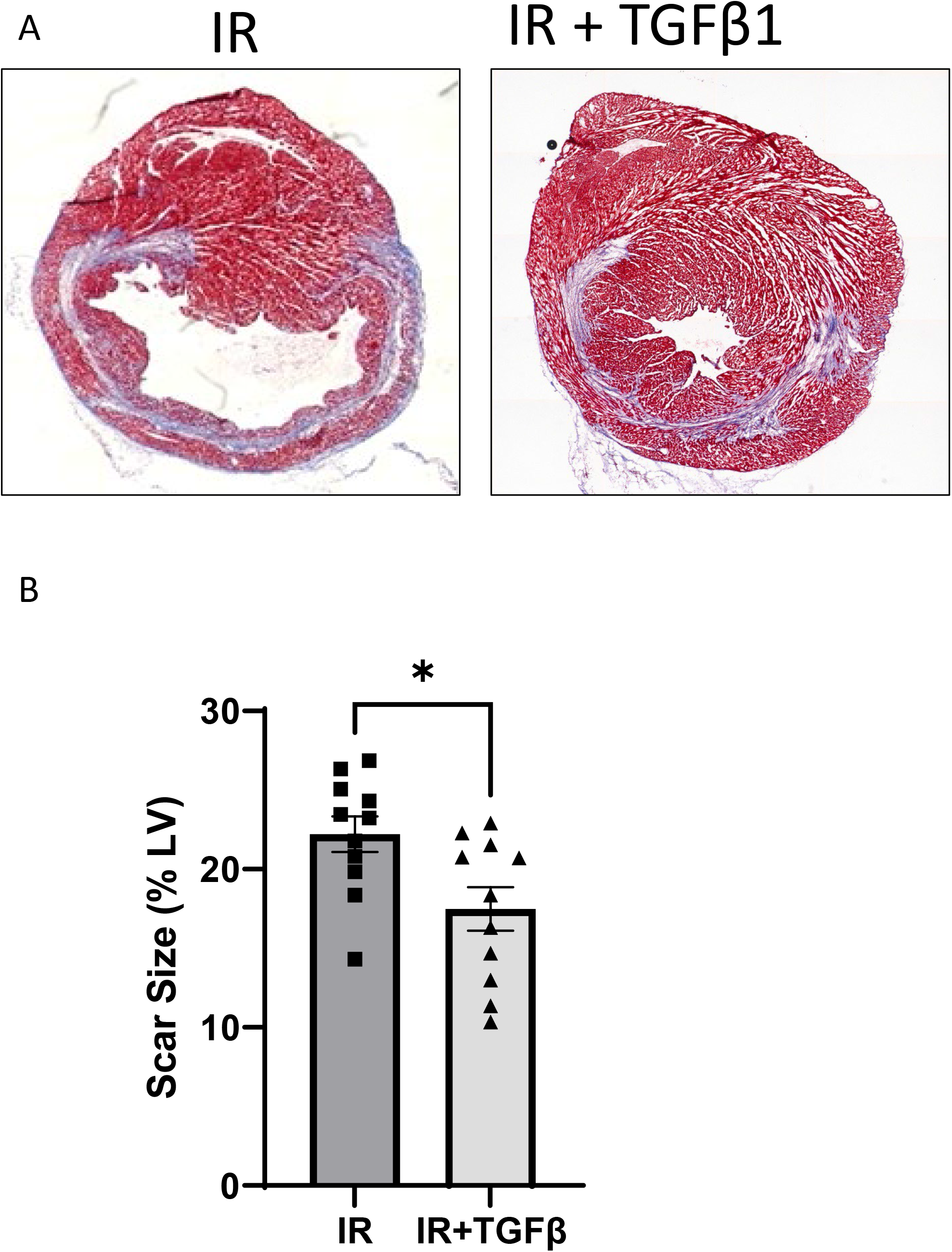
TGFβ1 treatment reduces scar size at 4 weeks post reperfusion. A: Masson’s trichrome stain was used to discriminate viable muscle (pink) from collagenous scar (blue). Analysis of >25 transverse sections through the heart was used to quantify scar size as percentage of left ventricle (%LV). B Scar size is significantly reduced in TGFβ1-treated mice compared with control mice subjected to the same cardiac injury of 60 minutes transient ischaemia (n=11/group). Data analysed by unpaired t test *p=0.015.

### Exogenous HpTGM reduces infarct size and inflammatory responses at 24h post reperfusion, and decreases mature scar size

Using the same mouse model of 60 minutes transient cardiac ischaemia, we first repeated the treatment schedule used above but with an equivalent dose of HpTGM in place of TGFβ1; that is, one dose via the tail vein immediately pre-infarction and a second dose via the same route immediately post-reperfusion. We found this treatment led to a very similar reduction of leukocyte infiltrate area and density in the heart at 24 hours post-reperfusion (Figure S3). This finding suggested that HpTGM had did indeed have similar anti-inflammatory properties to TGFβ1. However, any treatment commencing at the initiation of ischaemia is unlikely to be translatable to the clinic as this is prior to the time when most STEMI patients reach the hospital. We therefore decided to evaluate the effect of a single bolus of HpTGM given intravenously (via the tail vein) immediately (less than one minute) after reperfusion, to represent a clinically relevant time point for therapeutic delivery.

Control animals were subjected to the same cardiac injury and given saline, with the surgeon blinded to treatment. Single bolus HpTGM given at time of reperfusion led to a reduced mean infarct size (measured using CD45 infiltrate area) of 39%; p=0.038 and reduced mean leukocyte density by 44% (p<0.001) compared with saline treated animals (Figure 5A,B). HpTGM treatment also led to significantly reduced intra-cardiac expression of CCL2 (34% reduction p=0.0084) and IL1 β (49% reduction p=0.0036) compared with saline treated animals (Figure 5C,D) in a very similar way to exogenous TGFβ1 treatment. In addition, HpTGM appeared to be more effective than exogenous TGFβ1 in significantly reducing intracardiac expression of TNFα (30% reduction p=0.016) (Figure 5E). Endogenous *Tgfb1* transcripts are upregulated 2.5-fold in the injured heart at 24 hours following transient ischaemia, and this is not altered by HpTGM treatment (Figure 5F). Importantly, the single bolus of HpTGM given at time of reperfusion led to a significant reduction (28%; p=0.0076) of mature scar size at 4 weeks post-reperfusion (Figure 5G). As the early accruement of leukocytes in the infarcted heart tissue is mediated via leukocyte extravasation through the postcapillary venules we postulated that the TGFβ receptor in the vascular endothelium was required for the beneficial effect of HpTGM treatment. To test this we used a floxed *Tgfbr2* mouse carrying a tamoxifen-activated endothelial specific Cre (*Cdh5(BAC)-Cre-ert2)* that we have previously employed in developmental studies ^24, 35^(Figure 6). In agreement with the hypothesis, the effect of HpTGM on decreasing infarct size and reducing the density of leukocyte infiltrate is completely lost following *in vivo* deletion of the TGFβ receptor *Tgfbr2* specifically in endothelial cells. These data suggest this early benefit of delivering intravenous HpTGM in the acute setting was mediated via reduced leukocyte extravasation through the coronary vasculature.

**Figure 5.**
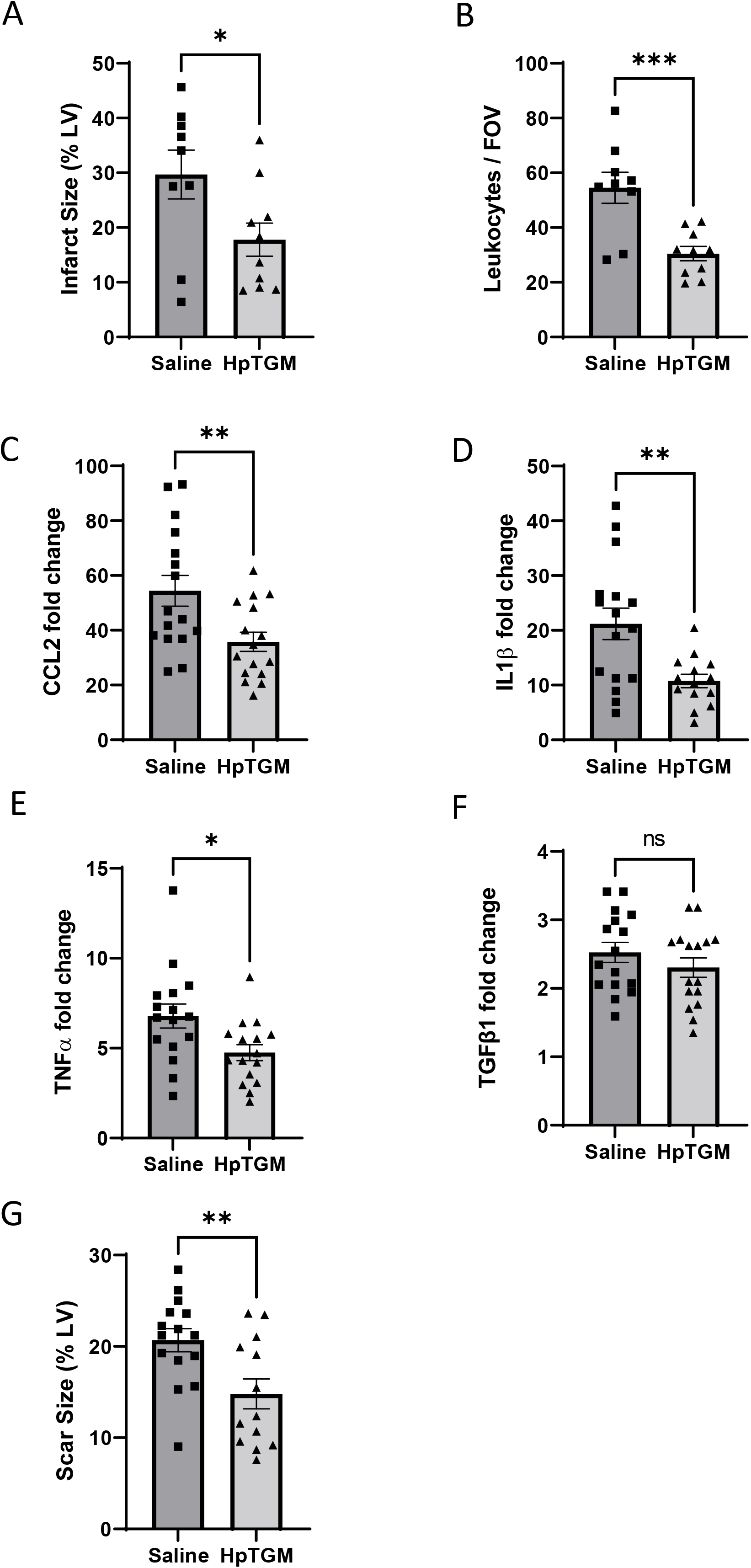
Treatment with exogenous HpTGM in a mouse model of myocardial infarction reduces infarct size, inflammatory cytokine expression and scar size. A Hearts were subject to 60 minutes infarction by transient occlusion of the LAD followed by reperfusion with or without HpTGM treatment (132ug/kg). Immunostaining of transverse heart sections with anti-CD45 (pan leucocyte marker) at 24 hours following reperfusion was used to measure the total area of infarct injury. HpTGM treatment leads to a significant reduction in the injured area. Mean values shown for 24 images per biological replicate. Data analysed by unpaired two tailed t test. *p<0.05 n=9-10/group B Anti-CD45 staining was used to measure the density of leukocyte accumulation in the injured region at 24hr reperfusion. HpTGM treatment leads to significantly reduced density of CD45+ immune cells in the injured area. Data analysed by unpaired two tailed t test. ***p<0.001 n=9-10/group. Abbreviations: lv left ventricle; rv right ventricle; FOV field of view. C,D,E,F : qPCR analysis of infarcted (60min) left ventricular heart tissue at 24h post reperfusion shows upregulation of cytokines CCL2, and TNFalpha in the left ventricle of IR hearts from saline treated mice compared with HpTGM treated mice. IL1b and TGFb1 transcript levels were similar in both groups. Gene expression data was normalised to housekeeping genes Hprt1 and Gapdh, and fold change in gene expression was calculated with respect to transcript levels of the target gene in naïve hearts. Data were analysed by unpaired two tailed t test. *p<0.05; **p<0.01. n=14-16/group. G: Scar size at 4 weeks post reperfusion is significantly reduced in HpTGM-treated mice compared with saline treated control mice subjected to the same cardiac injury of 60 minutes transient ligation of the LAD. Data was analysed by unpaired t test **p<0.01. n=13-15/group.

**Figure 6.**
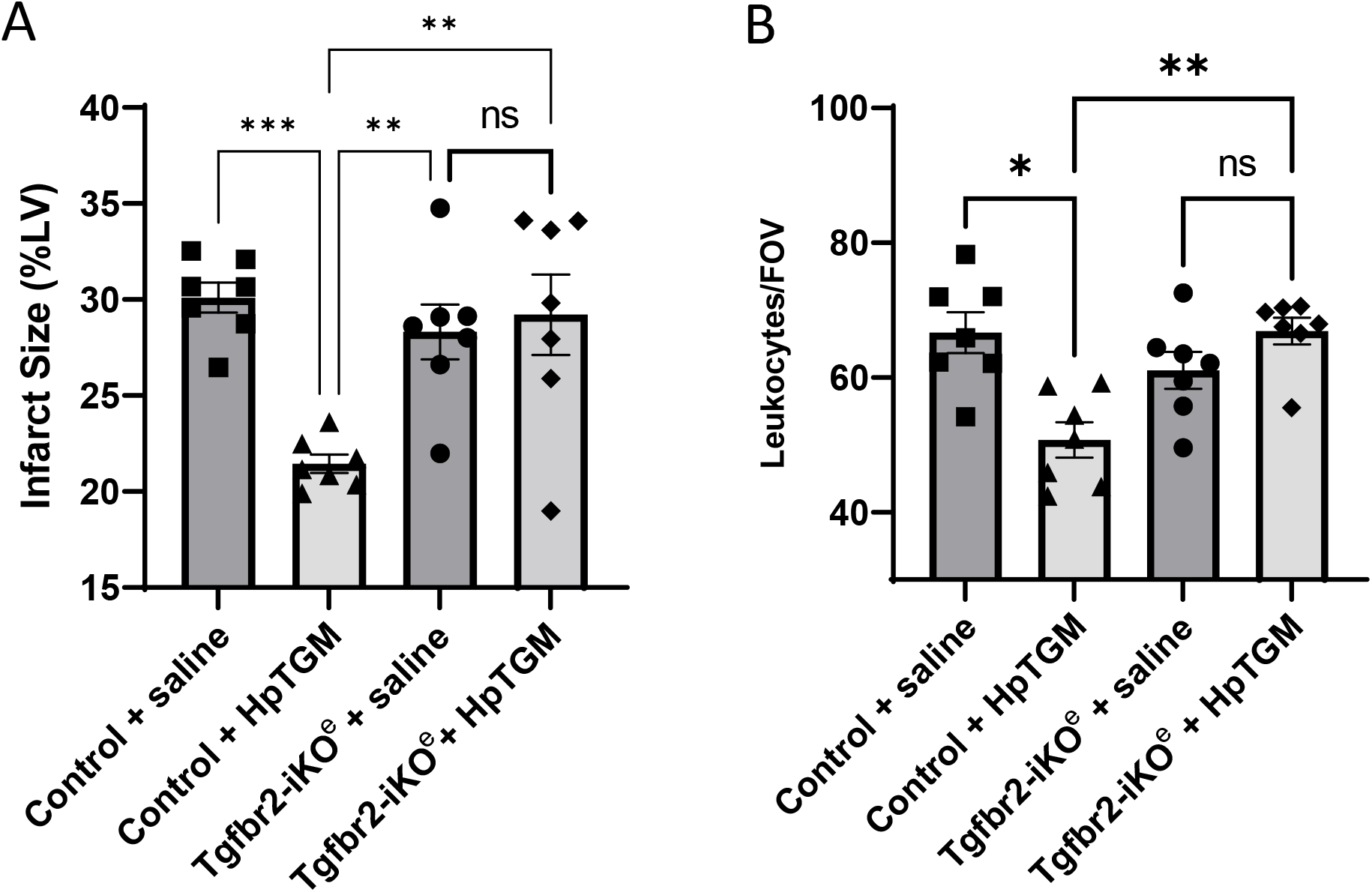
Cardiac protection of HpTGM treatment in the context of myocardial infarction is dependent on expression of TGFbeta receptor in endothelial cells. A,B Tgfbr2 floxed (Tgfbr2fl/fl) control mice and endothelial specific Tgfbr2 knockout (Tgfbr2-iKOe) mice were subject to 60 minutes occlusion of the LAD followed by reperfusion with or without HpTGM treatment (132ug/kg). Immunostaining of transverse heart sections with anti-CD45 (pan leucocyte marker) at 24 hours following reperfusion was used to measure the total area of infarct injury and the density of leukocytes per field of view (FOV). HpTGM treatment leads to a significant reduction in the injured area and density of leukocyte infiltrate in control Tgfbr2 fl/fl mice and this effect is lost in mice with no endothelial TGFBR2 receptor (Tgfbr2-iKOe). Data in A analysed by one way ANOVA with Tukey’s correction; data in B analysed by Kruskal-Wallis test with Dunn’s correction for multiple comparisons;. *p<0.05; ** p<0.01 n=7/group

## Discussion

Few clinical studies have examined circulating TGFβ levels in the acute STEMI setting. In stable coronary artery disease some interesting associations have been observed between low circulating TGF-β levels and poorer outcomes ^36^. Furthermore, patients with heart failure with preserved ejection fraction (HFpEF) have increased levels of circulating TGFβ1 compared with HF patients with reduced EF^37^. These are small hints that increased circulating levels of TGFβ1 might be beneficial in some cardiac patient groups. In the STEMI patients in this study, higher levels of circulating TGFβ1 at 24 hours post-PPCI were significantly associated with improved scar reduction after 3 months. A likely explanation of the beneficial effects of increased circulating TGFβ1 would relate to its potent anti-inflammatory properties. In order to initiate improved understanding of possible mechanisms we turned to a mouse model of transient myocardial ischaemia.

Numerous preclinical studies have pointed to the benefit of an anti-inflammatory approach in reducing infarct size. For example, targeting neutrophils directly or targeting IL1β has been shown to significantly reduce infarct size^38-42^. However, translating the benefits of anti-inflammatory therapy following MI to the clinic has proved challenging and led to doubts on the value of these animal studies. However, recent clinical findings have begun to change this perception and refocus energies on understanding the benefits of anti-inflammatory therapy in MI patients. For example, long term anti-inflammatory therapy targeting IL1β in MI patients with a pro-inflammatory blood profile (CANTOS trial) reduces the risk of recurrent cardiovascular events ^43^. Furthermore, the COLCOT trial revealed that early initiation of treatment post MI with the anti-inflammatory drug Colchicine was beneficial ^44^ and is consistent with our findings that early intervention is important to gain effective anti-inflammatory benefit. However, a simple and effective therapy at the time of PPCI to protect the remaining viable myocardium and thereby reduce the risk of progression to heart failure in STEMI patients remains an unmet clinical need.

This goal of dampening exuberant inflammatory responses also has to be reached within the context that some inflammation is beneficial and that timing of the therapeutic intervention is important. For example, early recruitment of myeloid cells to the injured heart tissue is required to remove dead cell debris. Also, following extravasation, monocytes differentiate to macrophages that within a few days make a key transition from so-called M1 to M2 macrophage to provide key reparative pro-angiogenic and pro-fibrogenic roles^45, 46^. In agreement with the anti-inflammatory protective effect of exogenous TGFβ1 in the acute setting, TGFβ <u>antagonists</u> delivered in the acute setting amplify the inflammatory response^47^. However, the same TGFβ antagonist treatment given later (3-7 days after MI) reduces adverse remodelling and myofibroblast apoptosis, consistent with a later detrimental role of TGFβ1^47-50^, that likely relates to its well-established role of promoting fibrosis^51^. *Thus TGFβ1 ligand therapy following MI is beneficial only when given within the very acute setting*.

As active TGFβ1 is very labile, with an immediate half-life of 11 mins and a terminal half-life of 60 mins^33^, delivery at the time of reperfusion means that the majority has cleared within 24 hours and well before it can influence pro-fibrotic intra-cardiac effects that normally initiate approximately 3 days following reperfusion in rodents (and slightly later in human). Furthermore, the absence of long term pro-fibrotic effects is consistent with the significantly reduced scar size we observe following exogenous TGFβ1 treatment in the acute setting. Nevertheless, to further reduce the risk of unwanted pro-fibrotic side effects, we turned to the TGFβ1 mimic HpTGM which has reduced pro-fibrotic properties^22^. We show that when delivered at the time of reperfusion, the anti-inflammatory and myocardial salvage effects of HpTGM are strikingly similar to those of TGFβ1. Furthermore, although HpTGM does not share molecular homology with TGFβ1 it signals through the same TGFβ receptor complex. Interestingly these early anti-inflammatory effects of HpTGM treatment appear to be mediated by the vascular endothelium as mice without endothelial TGFβ receptor 2 showed no detectable protective response to HpTGM compared with controls (Figure 6).

The importance of the vascular endothelium in myocardial ischaemic injury over and above the culprit occluded vessel(s) is widely recognised^52^. Here we show that TGFβ signalling in the vascular endothelium is required for mediating the early anti-inflammatory response to exogenous HpTGM treatment. This is consistent with the ability of TGFβ1 to reduce vascular endothelial expression of e-selectin, a protein that mediates leukocyte adhesion to the endothelium prior to extravasation^17, 18^. At 24 hours post reperfusion, neutrophils predominate the inflammatory infiltrate and TGFβ1 has been shown to inhibit transmigration of neutrophils through activated vascular endothelial cells^53^. Furthermore, intra-cardiac CCL2 plays a critical role in recruiting CCR2 expressing monocytes from the circulation and SMAD3 (a key transcription factor activated by the TGFβ receptor complex) mediates inhibition of CCL2 expression ^10^. Our study is unable to discriminate the cell specific source of the cytokines that are reduced following TGFβ1 or HpTGM treatment. However, their observed effects in reducing the numbers of leukocytes that have extravasated into the myocardium by 24 hours post-injury would inevitably have the corollary of reduced levels of leukocyte-expressed cytokines within the infarcted heart tissue.

Infarct size has a major impact on adverse cardiac remodelling, with larger infarcts leading over time to left ventricular dilatation, reduced end systolic volume and increased risk of heart failure. Thus, significant reductions in infarct size such as reported here can reduce the risk of progression to heart failure. The significant levels of cardiac protection by TGFβ1 and its mimic seen here are in stark contrast to our previous efforts to save infarcted myocardium by promoting angiogenesis. We generated substantial pro-angiogenic outcomes in the infarcted mouse heart using cardiosphere derived cell therapy but these cells failed to deliver long term benefit in heart function^27^. On the other hand, it is important to consider the pre-clinical model being used. Models of permanent myocardial infarction, where the ischaemic myocardium is almost entirely replaced by scar tissue leave little opportunity to rescue viable tissue. In contrast our current model of ischaemia reperfusion (as used in this study) represents a better preclinical model to evaluate therapies for STEMI patients undergoing PPCI. TGFβ1 has well established anti-inflammatory properties ^9-11^ whilst HpTGM is a parasitomimetic with great clinical potential. For example, recent work shows that delivery of HpTGM has a major anti-inflammatory effect in mouse models of colitis or airway inflammation ^54-56^. We show here that exogenous delivery of HpTGM at the time of coronary artery reperfusion dampens the pro-inflammatory response of coronary endothelial cells and reduces cardiac injury leading to increased myocardial salvage and reduced scar size with the corollary of improved prospects for long term cardiac function. These findings strongly support future work to further investigate the protective potential of HpTGM following cardiac infarction.

## Acknowledgements

This work was funded by the British Heart Foundation PG/18/57/33941 (HMA) and Wellcome Trust Awards 219530 and 104111(RMM)

